# Ex vivo human airway epithelia modelling reveals specific alterations in lung and hematopoietic stem cell transplant recipients

**DOI:** 10.64898/2026.06.15.732431

**Authors:** Louise Bondeelle, Junlu Sun, Sophie Clément, Claudio De Vito, Constant Gensous, Sheryline Loison, Yves Chalandon, Federica Giannotti, Gregory Berra, Romain Messe, Jérôme Le Goff, Jean Villard, Anne Bergeron, Caroline Tapparel

**Affiliations:** Department of Microbiology and Molecular Medicine, University of Geneva, Geneva, Switzerland; Department of Clinical Pathology, Geneva University Hospital, Geneva, Switzerland; Department of Haematology, Geneva University Hospital, Geneva, Switzerland; Pulmonology Department, Geneva University Hospitals, University of Geneva, Geneva, Switzerland; Université de Paris, AP-HP, Saint-Louis, laboratoire de virologie, 75000 Paris, France; Division of Transplantation Immunology, University Hospital of Geneva, Geneva, Switzerland; Université Paris Cité, UMR 1153 CRESS, ECSTRRA Team, F-75010, Paris, France

**Keywords:** bronchiolitis obliterans syndrome, hematopoietic stem cell transplantation, lung transplantation, chimerism, bronchial epithelium, lung allograft dysfunction, lung graft-versus-host disease

## Abstract

Deterioration of lung function is a major cause of long-term morbidity after hematopoietic stem cell transplantation (HSCT) and lung transplantation (LT). In both settings, obliterative bronchiolitis represents the most common final pathway, with bronchiolitis obliterans syndrome (BOS), representing its clinical correlate. Understanding of the pathophysiological mechanisms leading to BOS is limited by restricted access to human lung tissue and the imperfect relevance of animal models. We hypothesize that transplantation procedures cause bronchial epithelial damage that promotes the development of BOS. To investigate this, we established *ex vivo* human airway epithelia (HAE) cultures from bronchial biopsies of HSCT and LT recipients, collected prior to the development of BOS, and compared them with non-transplant controls.

HAE from HSCT recipients exhibited reduced tissue differentiation ability, associated with defect in mucociliary clearance and impaired barrier integrity, most markedly in one patient who subsequently developed BOS. In contrast, LT-derived HAE showed normal mucociliary clearance and barrier integrity but displayed increased mucin secretion. Donor and recipient-derived cells were detected in both paraffin-embedded biopsies and reconstructed HAE derived from transplant recipients, demonstrating epithelial chimerism. Our data highlight specific modifications of the airway epithelium after LT and HSCT that may represent a first trigger for subsequent BOS development.

## Introduction

Progressive impairment of pulmonary function is a major determinant of long-term morbidity and mortality following both allogeneic hematopoietic stem cell transplantation (HSCT) and lung transplantation (LT)^1^. Despite distinct clinical settings, both LT and HSCT can be complicated by obliterative bronchiolitis (OB). OB is histologically characterized by an inflammatory infiltration of small airways, and fibrotic remodelling leading to luminal obliteration^1^. As transbronchial biopsies have a low diagnostic yield and surgical biopsies are considered too invasive, histologic confirmation is seldom obtained. Consequently, the diagnosis relies on respiratory function criteria and is referred to as bronchiolitis obliterans syndrome (BOS). BOS is defined as a sustained ≥10% decline in Forced Expiratory Volume in 1 second (FEV1) from baseline, together with an obstructive ventilatory defect^2,3^. While some patients with a 10% FEV1 decline recover without developing fixed airway obstruction, a ≥ 10% decrease in FEV1 from baseline is a predictor of BOS^1,2,4^.

OB/BOS is considered an alloimmune complication, with increasing evidence suggesting that bronchial epithelial insults associated with both HSCT and LT procedures, may act as initiating triggers^1^. The bronchial epithelium is a dynamic regulator of airway integrity, repair and immune defence. It comprises multiple specialized cell populations, including basal, supra-basal and club cells, with progenitor capacity, as well as ciliated and secretory lineages that ensure mucociliary clearance, barrier integrity and antimicrobial defence^5–7^. Disruption of these epithelial functions weakens both the physical and immune barriers, compromising airway repair and promoting aberrant remodelling that culminates in OB^8–10^. In LT, ischemia-reperfusion perioperative stress can impact epithelial integrity^11^. In HSCT, pre-transplant conditioning regimens are also known to affect epithelial compartments, as shown in other tissues and experimental models^12–14^. In addition, post-transplant complications such as infections or treatment-related toxicities may further influence epithelial structure and function^1^. Despite these observations, the extent and nature of epithelial alterations in OB, and their variability across patients and transplant settings, remain unclear.

Chimeric bronchial epithelial cells have been reported in both LT and HSCT, consistent with epithelial cell replacement after injury^15,16^. Co-existence of donor- and recipient-derived epithelial cells may influence repair, immune recognition and tolerance, shaping airway remodelling. However, the functional consequences of epithelial chimerism in OB pathogenesis is unknown^15,16^.

Although existing animal models reproduce some immune or fibrotic features, they only partially recapitulate the chronic, progressive obliteration of the small airways in humans^17^. Anatomically, murine airways differ from human airways with fewer conducting airway generations, lack submucosal glands in distal bronchi and a more uniform epithelial architecture^18^. Functionally, murine airway repair is dominated by rapid epithelial restitution, lineage plasticity and efficient progenitor-driven regeneration with limited persistence of fibroproliferative remodelling, in contrast to the delayed, dysregulated repair and sustained fibrosis characteristic of human OB^19^. These limitations are addressed using human airway epithelia (HAE), which are reconstituted *ex vivo* from patient bronchial biopsies and preserve the main epithelial lineages and disease-associated phenotype^17^. Used in asthma and infection studies, their relevance to HSCT and LT associated airway disease has yet to be established. Nonetheless, HAE cultures provide a physiologically relevant model to investigate epithelial differentiation, mucociliary function, and responses to environmental, infectious, or pharmacological challenges in the setting of transplantation.

We hypothesized that post-transplant OB/BOS result from alterations in airway epithelial integrity, triggered by repeated transplant-related insults. To test this, we used in-house *ex vivo* HAE models reconstituted from bronchial biopsies obtained from HSCT and LT recipients, and integrated analyses. We compared these models to HAE derived from non-transplant (NT) controls and with the corresponding original paraffin-embedded patient biopsies. By studying HSCT and LT in parallel, we aimed to identify common epithelial defects that could be associated with BOS development, as well as context-specific mechanisms t unique to either HSCT or LT.

We found consistent epithelial abnormalities. LT-derived epithelia showed increased mucin secretion, while HSCT-derived epithelia showed impaired differentiation, reduced mucociliary clearance and compromised barrier integrity. Epithelial chimerism observed in FFPE biopsies was also detected in reconstituted HAE from both HSCT and LT, demonstrating faithfull recapitulation *in vitro,* providing a platform for functional investigations.

## Materials and Methods

### Development of ex-vivo reconstituted HAE cultured at an air-liquid interface

Transplant recipients (LT and HSCT) and NT undergoing bronchoscopy at Geneva University Hospitals who met the inclusion criteria and provided their informed consent were enrolled in the study (Table 1).

**Table 1.**
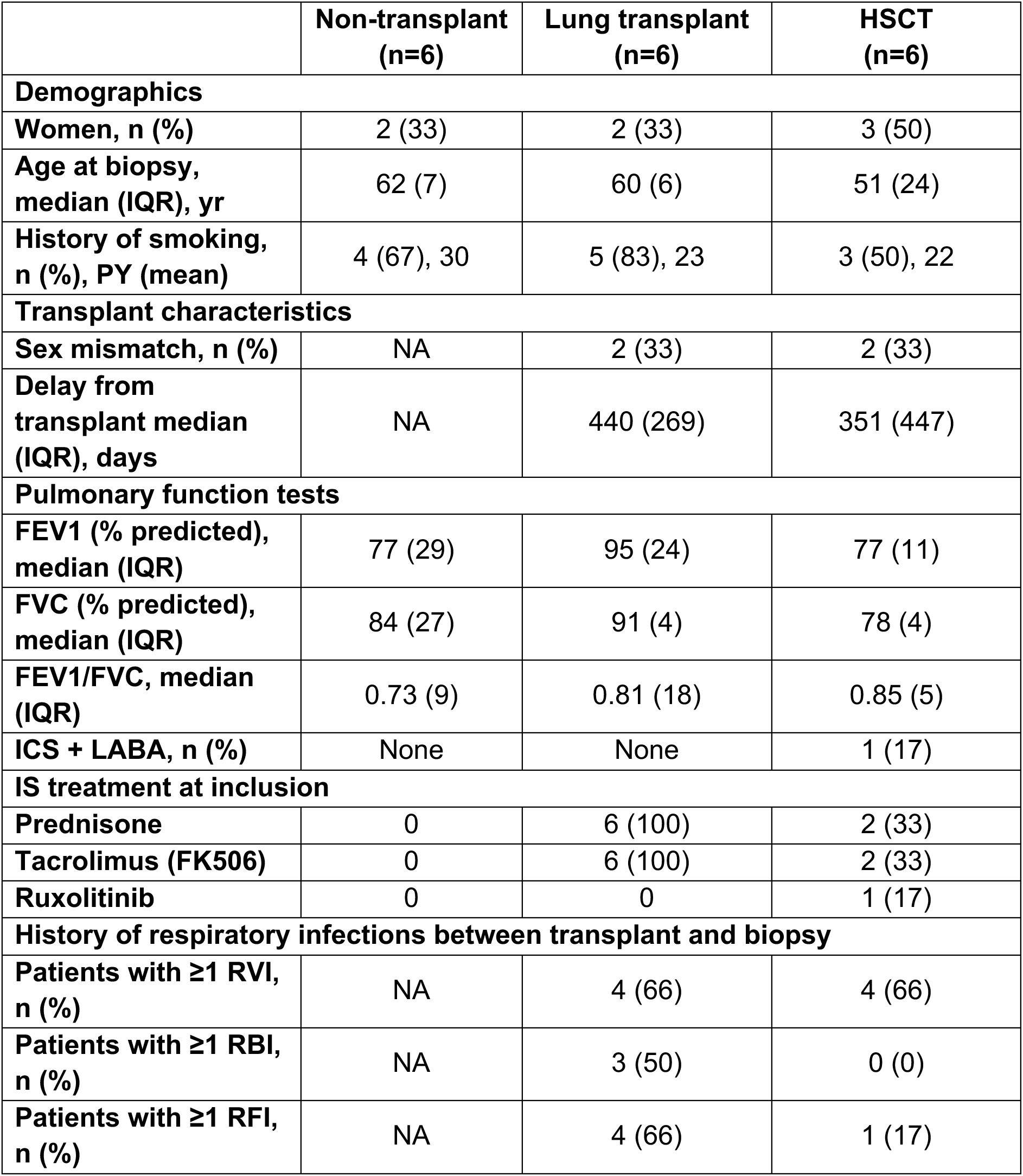
Patient characteristics. IQR: interquartile (Q1-Q3), yr: year. PY: pack-year (number of cigarette packs per day x the number of years of smoking), FEV1: forced expiratory volume in 1 second, FVC: forced vital capacity, ICS: inhaled corticosteroid, LABA: long-acting beta2 agonists, IS: RVI; respiratory viral infection (HPIV3: human parainfluenza virus type 3; RHV: rhinovirus; RSV: respiratory syncytial virus; MPV: metapneumovirus; CoroOC43: coronavirus OC43); RBI: respiratory bacterial infection, RFI: respiratory fungal infection, NA: not applicable. See Supplementary Tables S1-3 for individual patient data.

The inclusion criteria were as follows: (i) for NT controls, bronchoscopy performed for the investigation of a lung nodule or a benign disease, with no history of chemotherapy or radiotherapy; (ii) for LT recipients, bronchoscopy performed during routine follow-up (up to 5 years post-LT); and (iii) for HSCT recipients, bronchoscopy performed in the context of a recent decline in FEV1 exceeding 10% from baseline in HSCT (up to 3 years post-HSCT).

None of the transplant recipients had BOS criteria at the time of the biopsy. Patients with a respiratory infection were excluded. Among the enrolled subjects, only sample that yielded successful HAE reconstitution were included in the final analyses. The study was approved by Swiss ethics N°2023-00140).

### Bronchial Biopsy & Cell Culture

Flexible bronchoscopy was performed, and three airway mucosal biopsy specimens were obtained from each patient as distally as possible within the bronchial tree for *ex vivo* culture. Respiratory samples obtained during the bronchoscopy were processed for standard bacterial and fungal cultures, and for multiplex viral PCR (Tianlong® 12-virus kit) targeting influenza A/B, rhinovirus, respiratory syncytial virus, adenovirus, metapneumovirus, coronaviruses and parainfluenza viruses. Biopsies were excluded from the study when respiratory samples tested positive for infection (Fig. 1). After expansion, primary bronchial epithelial cells were cultured under air-liquid interface (ALI) conditions, differentiating into mucociliary phenotypes within 28 days as previously described^20^.

**Figure 1.**
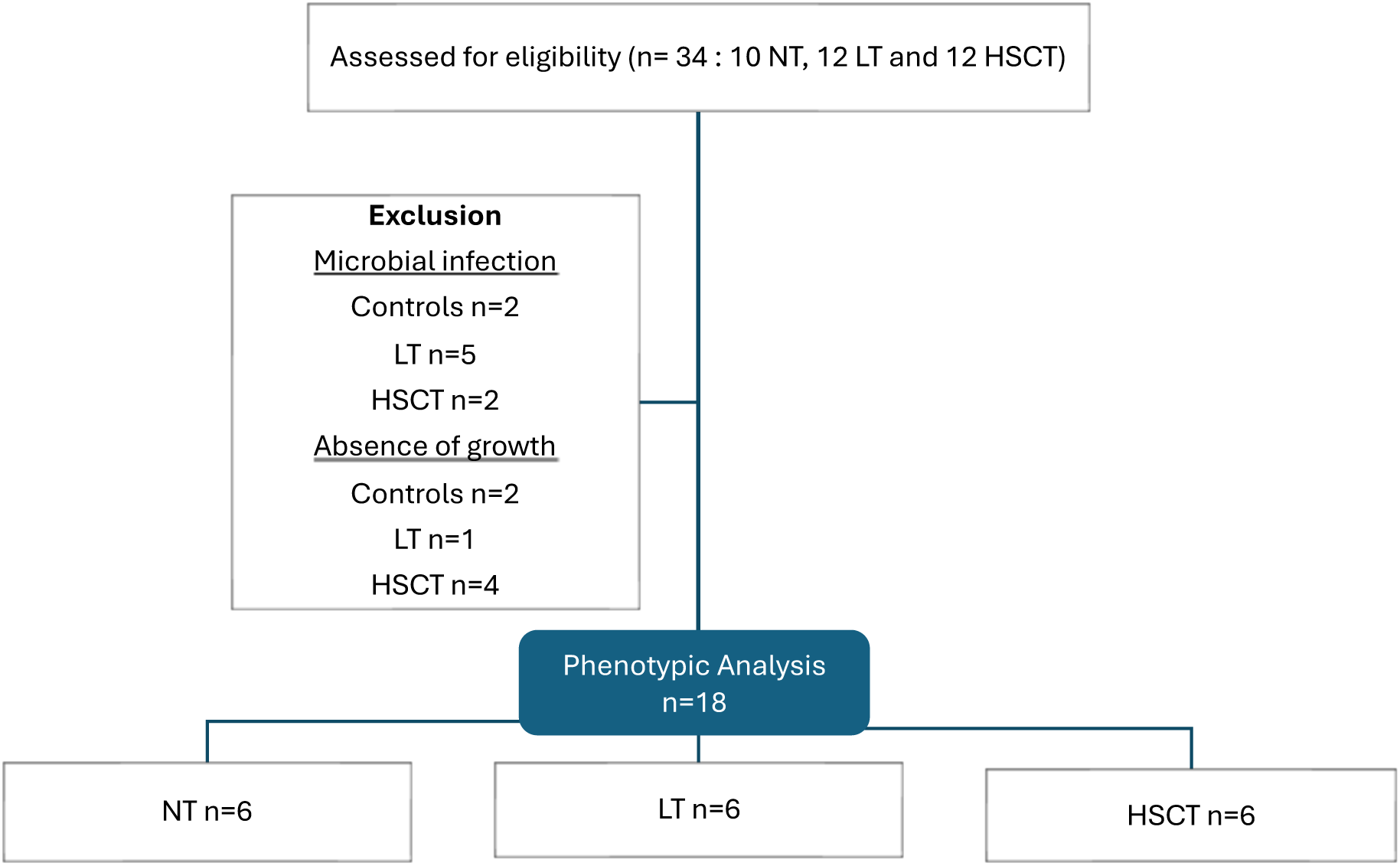
Study workflow for HAE reconstitution and phenotypic analysis. Bronchial biopsies were obtained from 34 patients and assessed for eligibility. Samples were excluded if presence of a pathogen or when HAE growth failed. Fully reconstituted HAE were successfully generated for 18 patients, with six samples in each group: non-transplanted (NT), lung transplant (LT) and hematopoietic stem cell transplant (HSCT). Phenotypic analyses, including morphological, immunofluorescence and functional characterization were performed on these reconstituted HAE.

### Histological & Immunofluorescence Analysis

Routine biopsies collected during the same bronchoscopy were paraffin-embedded (FFPE) and stained with haematoxylin and eosin (H&E). Except for FISH (see below), no additional staining was performed on these sections, which were taken for analysis as part of routine care. These samples served as a reference for comparison with the HAE cultures. Fully differentiated HAE cultures were fixed with 4% paraformaldehyde, permeabilized with fix and perm buffer (ThermoFischer, 4222-26, 1/10) and immunostained with primary antibodies against β-tubulin IV (ciliated cells, Abcam 179504, 1/200), MUC5AC (mucous cells, conjugated, Abcam 3649, 1/200) and cytokeratin 5 (basal cells, conjugated, Abcam EP1601Y, 1/500). Alexa fluor-conjugated secondary antibodies (Alexa fluor 488 1/200) were applied, nuclei were counterstained with DAPI, and membranes mounted in antifade medium. Imaging and Z-stack reconstruction were performed by confocal microscopy (Axio Imager, Z2 LSM800) using Zen software.

### Mucus & Mucociliary Clearance (MCC) analysis

Mucin secretion was quantified via Enzyme-linked Lectin Assay (ELLA) (Sigma-Aldrich, M2378) according to the manufacturer’s instructions. MCC was monitored using polystyrene microbeads (Sigma-Aldrich 84135) captured at 2 fps using a Sony XCD-U100CR camera, and beads velocity was analysed with ImageProPlus 6.0.

### Trans-Epithelial Electrical Resistance Measurement

TEER (Ω·cm²) was measured with a Millicell ERS volt-ohm meter after one month under ALI conditions, after subtracting blank values, as previously described^21^.

### Quantitative PCR analysis of gene expression

Samples were lysed in TRK lysis buffer (E.Z.N.A. RNA extraction Kit; Omega Bio-Tekand) and total RNA was extracted according to the manufacturer’s recommendations. Gene expression was quantified by TaqMan-based one step reverse transcription quantitative PCR (RT-qPCR) (ThermoFisher Scientific, Cat. No. 4444556). We selected specific TaqMan primers and probes to assess mRNA expression of basal cells markers (KRT5, Hs00361185_m1; TP63, Hs00978343_m1) ciliated cells markers (FOXJ1, Hs00230964_m1; DNALI1, Hs00185750_m1), goblet markers (MUC5AC, Hs01365616_m1; TFF3, Hs00902278_m1), the club cell marker (SCGB1A1 HS00171092_m1) and the proliferating cell marker (MKI67, Mm01278617_m1). GAPDH (Assay ID 4310884E) was used as the housekeeping gene. All assays were obtained from ThermoFisher Scientific. Amplification was performed with the following cycling conditions: 30 min at 50°C, 15 min at 95°C, followed by 45 cycles of 15s at 94°C and 1min at 60°C (7500 Fast Real-time PCR system, Applied Biosystems). Relative gene expression was calculated using the comparative Ct (ΔΔCt) method, using non-transplant as the reference condition.

### Detection of epithelial chimerism in original patient biopsies and corresponding reconstituted HAE derived from HSCT and LT recipients

Airway epithelial chimerism was evaluated in FFPE biopsies and reconstituted HAE from sex-mismatched transplant recipients using FISH with XY probes. In HSCT recipients, chimerism was further quantified with AlloSeq 2.2.0 HCT, a targeted next-generation sequencing assay enabling donor-recipient DNA discrimination. For detailed protocol see supplementary methods.

### Statistics

Statistical analyses were performed using GraphPad Prism (version 10). Pairwise comparisons between predefined groups (NT vs LT, NT vs HSCT and LT vs HSCT) were performed using two-sided non-parametric Mann-Whitney tests. P values <0.05 were considered statistically significant.

## Results

### Patient characteristics

Bronchial biopsy specimens obtained from 18 subjects (6 NT, 6 HSCT and 6 LT) were included in the study. Their clinical characteristics are given in Table 1 while details for each subject are provided in the <u>supplementary material</u> (Tables S1-3).

### Reconstitution and morphological characterization of reconstituted HAE

ALI-cultured HAE from LT, HSCT and NT subjects (see study workflow in Fig. 1) were generated from patient bronchial biopsies as schematized in Fig. 2A. Thirty-four subjects (10 NT, 12 HSCT and 12 LT recipients) were initially included but tissues were successfully reconstructed for 18 of them (6 NT, 6 HSCT and 6 LT recipients).

**Figure 2.**
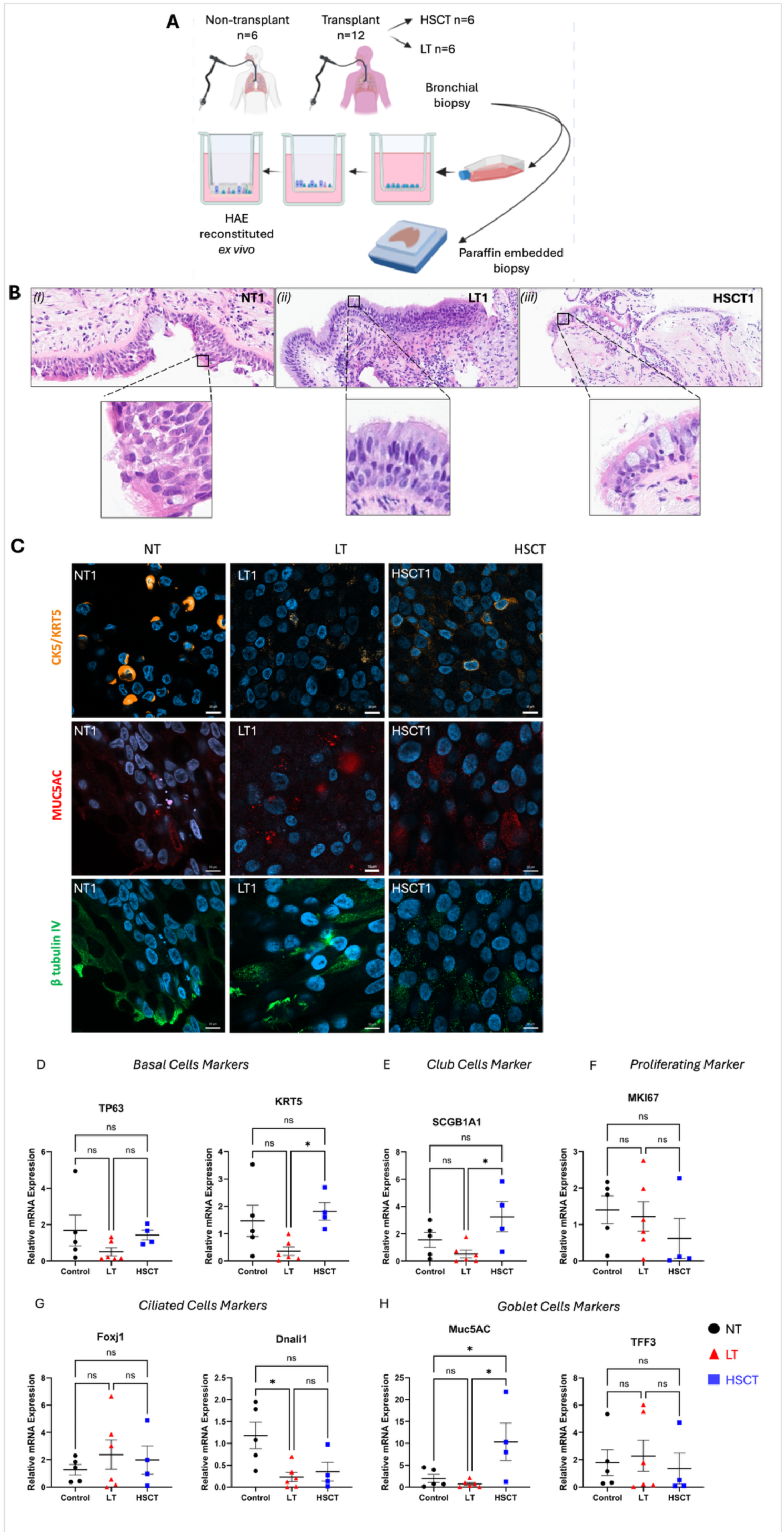
Morphological characterization of bronchial biopsies and reconstituted HAE. (A) Bronchial biopsies obtained by bronchoscopy from each patient were used both to reconstitute an airway epithelium under air-liquid interface conditions and to be paraffin-embedded for histological comparison of bronchial epithelium. (B) Haematoxylin-eosin (HE) staining of paraffin-embedded sections prepared from bronchial biopsies of NT (i), LT (ii), and HSCT (iii); shown at 10x and 40x magnification. (C) Immunofluorescence of reconstituted HAE, highlighting basal cells (CK5, orange), goblet cells (MUC5AC, red) and ciliated cells (β-tubulin IV, green). Representative images from HAE derived from one subject in each group are shown (see Fig. S1 for all donors). Confocal microscopy x63 objective, scale bars: 10μm. Panels D-H show mRNA expression levels of marker genes for specific epithelial cell types: basal cells (TP63, KRT5, panel D), club cells (SCGB1A1, panel E), proliferating cells (MKi67, panel F), ciliated cells (FOXJ1, DNALI1, panel G) and goblet cells (MUC5AC, TFF3, panel H). Analyses were performed on mRNA isolated from HAE cultured at ALI conditions for 28 days. Significance was determined using the non-parametric Mann-Whitney test. Data are presented as median ± IQR of the HAE for each subject (mean of technical duplicates). *p<0.05, **p<0.01. HAE: human airway epithelium, LT: lung transplant, HSCT: hematopoietic stem cell transplant.

During the reconstruction phase, successful HAE reconstruction was initially achieved in 8/12 HSCT (66%), in 8/10 NT (80%) and in 11/12 LT (92%) samples. Two NT cultures, two HSCT cultures and 5 LT cultures were subsequently excluded because of contamination, resulting in a final analytical cohort of six HAE cultures per group. The lower reconstruction success rate observed in HSCT-derived HAE may suggest impaired regenerative capacity of epithelial progenitor cells following HSCT. Reconstituted epithelia were subsequently analysed for morphological differences and compared with the corresponding native epithelium from FFPE bronchial biopsies. Images of HAE immunostaining with cell-type-specific markers and of haematoxylin-eosin-stained FFPE biopsies are shown in Fig. S1 and S2, respectively, for all donors, while Fig. 2B presents illustrative exanples from one donor per condition.

After reconstruction, the main epithelial cell types were detected in all tissues, with variable proportion across the different samples. In LT and NT-derived HAE, all cell types were present at broadly similar levels, although a slight decrease in basal cell maker staining was observed in LT samples (Fig.S1). In HSCT-derived HAE, variability across samples was observed: three out of six (HSCT 1-3) showed a marked reduction in β-tubulin IV–positive ciliated cells, whereas the extent of MUC5AC-positive goblet cells was more heterogeneous, with HSCT1 showing levels comparable to some NT-derived samples (Fig. S1). Histological analysis of FFPE biopsies collected from the same patients (Fig. S2) also highlighted a well-differentiated epithelium with preserved ciliated morphology in NT samples, in most LT samples and in four of six HSCT samples. By contrast, 2 HSCT recipients (HSCT2 and HSCT3) exhibited a pronounced loss of ciliated and goblet cells in FFPE biopsies with metaplastic changes consistent with defective epithelial differentiation observed in HAE from the same patients and with the overall decreased reconstruction success rate of HAE from HSCT donors.

To better characterize the differentiation within each HAE group, differentiated tissues were lysed and RNA was extracted for RT-qPCR analysis of cell-specific markers. Markers of basal (KRT5, TP63), club (SCGB1A1), goblet (MUC5AC, TFF3), ciliated (FOXJ1 and DNALI1) and proliferating cells (MKi67) (Fig. 2D-H) were quantified and normalised to GAPDH.

Due to the limited sample size and the substantial inter-individual variability, only a few differences reached statistical significance. DNALI1 expression was significantly reduced in both LT- and HSCT-derived HAE, while KRT5 expression was significantly increased in HSCT-derived HAE compared with LT-derived tissues.

Several markers showed group trends, with lower expression of progenitor (KRT5, TP63 and SCGB1A1) and goblet-associated genes in LT-derived HAE, and higher expression in HSCT-derived HAE, while an opposite trend was observed for MKI67, a proliferative marker.

### Functional characterization of reconstituted HAE derived from LT, HSCT and NT

All reconstituted HAE were subjected to functional analyses. Barrier integrity, mucociliary clearance and mucin secretion were evaluated using respectively TEER, MCC and ELLA assays. Mucociliary clearance and barrier integrity were significantly decreased in HSCT-derived HAE. Mucus secretion was significantly increased in LT-derived HAE compared with NT, wheres no significant difference was observed between HSCT-derived HAE and NT.

### Link between epithelial phenotype and clinical parameters

Variability in epithelial phenotypes was analysed in the context of clinical parameters including post-transplant respiratory viral infections (RVI), treatment history and clinical evolution.

Recipients without documented RVI after transplantation (HSCT1, HSCT6, LT4 and LT5; tables S2 and S3) displayed variable epithelial features. Among them, HSCT6, LT4 and LT5 exhibited well-differentiated airway epithelia and were characterized by preserved TEER, efficient MCC and the highest mucin secretion (tables S4-6). In contrast, recipients with three or more documented RVI (LT1, LT2, LT3, LT6, HSCT2, HSCT3 and HSCT5) exhibited heterogeneous epithelial alterations. Among LT recipients, only LT2 showed a clear reduction in mucin secretion and MCC, while TEER remained within the normal range and was not consistently reduced in the other LT recipients. In HSCT, epithelial dysfunction was more evident, particularly in HSCT2 and HSCT3, with milder impairment in HSCT5. HSCT3-derived HAE displayed the most severe epithelial barrier phenotype with the lowest TEER (252 Ω·cm² *versus* HSCT group median 518 Ω·cm² [IQR 41]), markedly reduced MCC (5,79 µm/s *versus* 20.6 µm/s, [IQR 23.6]) and decreased mucin secretion (0.2765 OD, *versus* 0.4373 OD, [IQR 0.13836]) (Fig. 3A-C). Strikingly, the corresponding paraffin embedded biopsy also presented a metaplastic epithelium (Fig. S2). Clinical follow up highlighted development of BOS three months after the biopsy was performed (Fig. S3). Notably, this patient exhibited a sustained decline in pulmonary function, with FEV1 and FVC falling below 75% of baseline shortly after transplantation and progressively worsening over time. This coourse is consistent with impaired epithelial differentiation observed in both the native biopsy and the reconstructed HAE.

**Figure 3.**
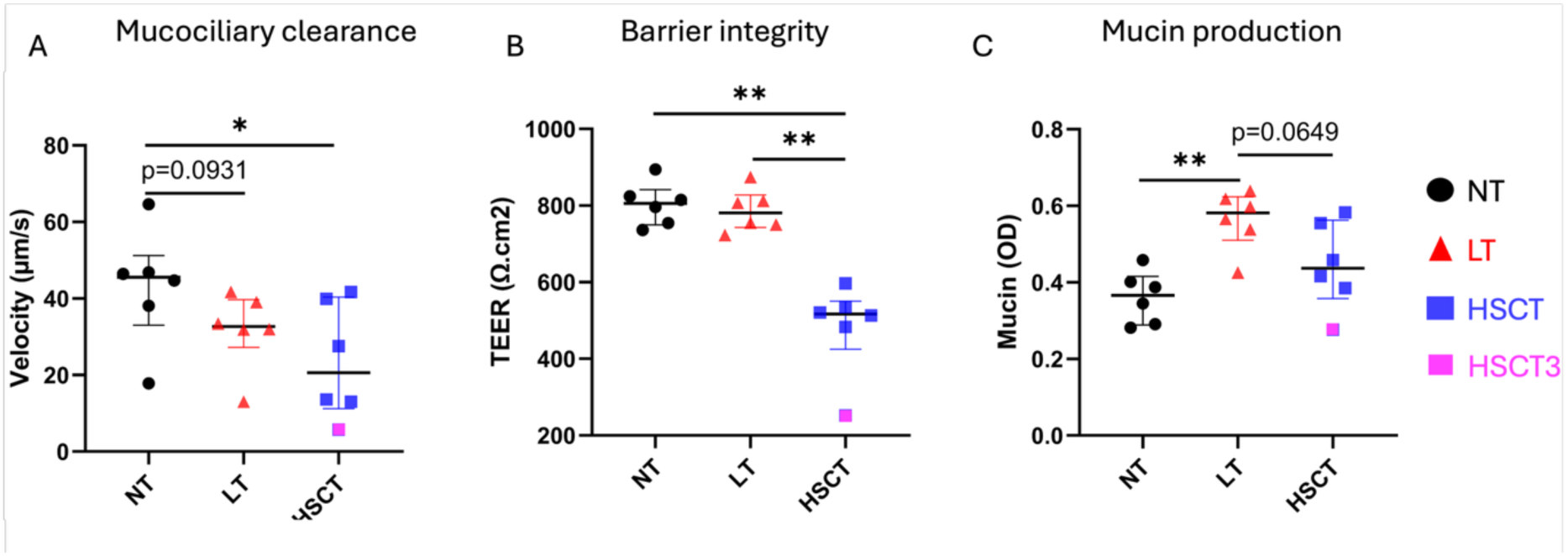
Functional assessment of reconstituted human airway epithelia. (A) <u>Mucociliary clearance</u>, quantified as microbeads velocity across HAE apical surface, is markedly reduced in HSCT-derived HAEs (median 20.6 ms^-1^; IQR 23.6) compared with NT-derived HAE (median 45.6ms-1; IQR 6.9) and LT-derived HAE (median 32.7 ms^-1^; IQR 5.7). (B) <u>Barrier integrity</u>, assessed by transepithelial electrical resistance is compromised in HSCT-derived HAE (median 517 Ω.cm^2^; IQR 40), compared to NT (median 805 Ω.cm^2^; IQR 58) and LT (median 781 Ω.cm^2^; IQR 59). (C) <u>Mucin secretion</u>, quantified by ELLA assay, is significantly increased in LT-derived HAE compared with NT (LT; median OD 0.58156; IQR 0.06787; NT: OD 0.3663, IQR 0.09358; HSCT OD 0.4373, IQR 0.13836). HSCT3 is highlighted in pink. Significance was determined using the non-parametric Mann-Whitney test. Data are presented as median ± IQR. * p<0.05, **p<0.01. TEER measurements were obtained from independent biological triplicates, while mucin and MCC analyses were performed using independent biological duplicates.

### Detection of epithelial chimerism in biopsies and reconstituted HAE from transplant recipients

Epithelial chimerism was assessed in both FFPE bronchial biopsies and HAE reconstituted from LT and HSCT recipients with sex-mismatched donors (HSCT1, HSCT4, LT2 and LT4). In addition, HSCT3, in which both donor and recipient were of the same sex (XX), was included as control (Tables S2, S3). In HSCT-derived HAE, the airway epithelial compartment is expected to originate from the transplant recipient, while in LT-derived HAE, the airway epithelial cells are expected to originate from the lung donor.

Using RNA fluorescence in situ hybridisation (RNA-FISH) with X- and Y-chromosome probes, epithelial cells of donor or recipient origin were identified in all sex-mismatched recipients (Fig. 4A). As expected, only XX cells were detected in the sex-matched control HSCT3. Cells of donor origin were identified in HSCT-derived HAE, corresponding to female (XX) cells in male recipients HSCT1 and HSCT4, while cells of recipient origin were detected in LT-derived HAE, corresponding to female (XX) cells in the male recipient LT2 and male (XY) cells in the female recipient LT4.

**Figure 4.**
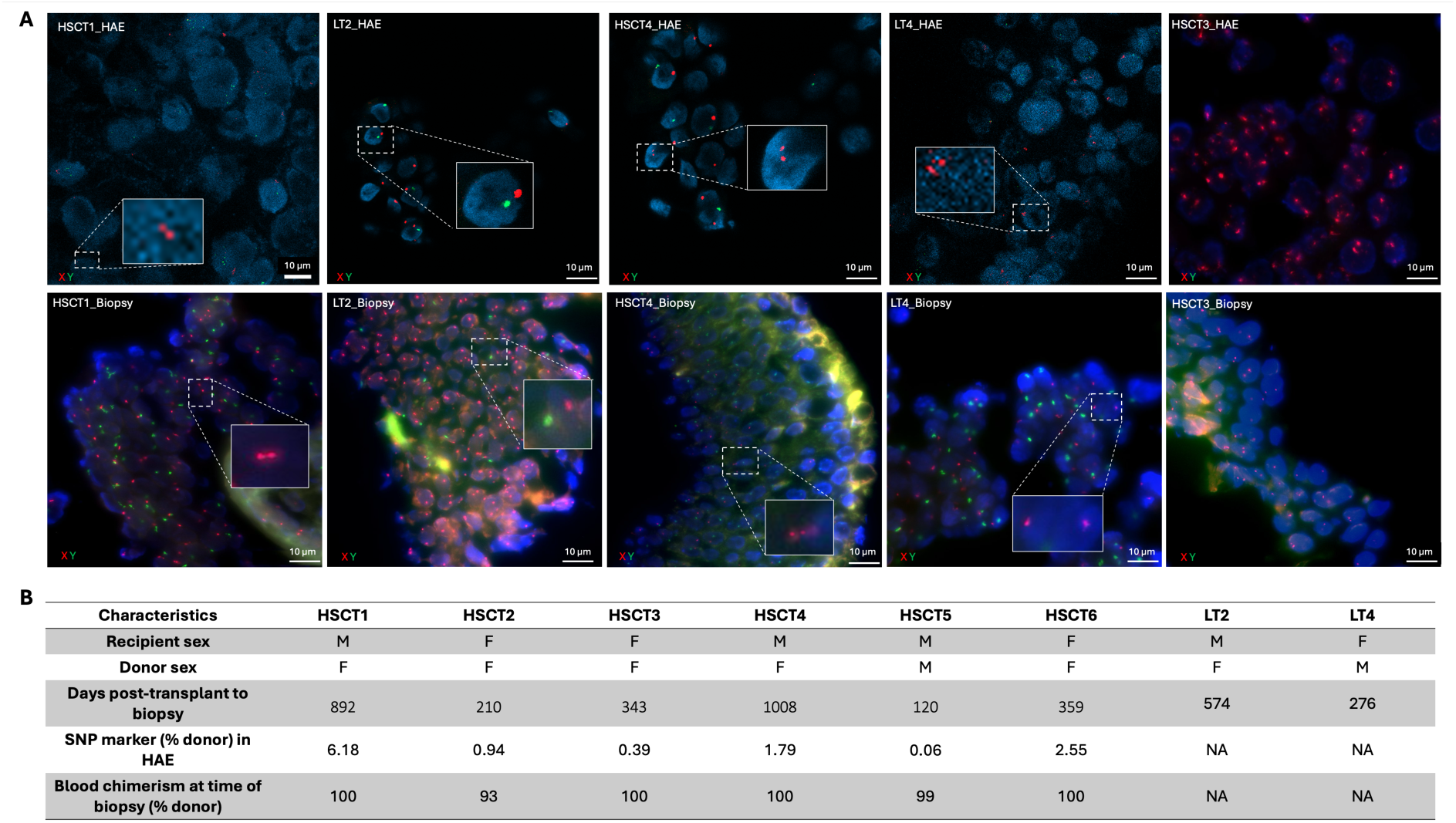
Chimerism in HAE: evidence from FISH in HSCT and LT recipients and from NGS in HSCT. (A) Airway epithelial chimerism detected by FISH in cells dissociated from reconstituted HAE (HSCT1, HSCT4, LT2 and LT4) and in the corresponding FFPE bronchial biopsies. The highlighted chimeric cells were rare in the whole tissue. LT2, HSCT1 and HSCT4 were male recipients with female donors, whereas LT4 was a female recipient with a male donor. HSCT3 was a female recipient with female donor and serves as a negative control demonstrating the absence of false Y-positive. Nuclei are counterstained with DAPI (blue), X chromosomes in red and Y chromosomes in green. Chimeric and non-chimeric cells are highlighted by white dashed squares. Confocal microscopy, x40 objective, scale bars: 10μm. (B) Characteristics of HSCT and recipients with corresponding levels of epithelial chimerism in reconstituted HAE and blood chimerism, determined by SNP-based NGS.

To complement these qualitative observations and estimate the extent of epithelial chimerism, SNP-based next generation sequencing (NGS) was performed in HSCT cases for which donor genomic sequences were available to detect and quantify chimerism in reconstituted HAE. This approach could not be applied to the original HSCT biopsies because the presence of blood cells would have biased the interpretation of the results. Donor-derived epithelial signatures were consistently detected, although at levels ranging from <1% to 6% (Fig. 4B).

## Discussion

In this study, we hypothesized that transplantation-related procedures induce bronchial epithelial alterations that may contribute to the development of bronchiolitis obliterans which is a common complication of both HSCT and LT. By establishing *ex vivo* HAE from bronchial biopsies collected from HSCT and LT recipients and comparing them with NT-derived HAE controls, we aimed to identify common or specific epithelial features across transplant settings consistent with the pathophysiology of OB.

Our results revealed two distinct epithelial phenotypes. LT-derived epithelia were comparable to NT-derived HAE, whereas HSCT-derived epithelia displayed more heterogeneous and frequently altered features. These findings suggest that despite a shared alloimmune context, epithelial remodelling differs between LT and HSCT, indicating that factors beyond immunity contribute to epithelial alterations and that OB may develop through distinct mechanisms.

### Epithelial phenotypes and heterogeneity

#### HSCT epithelia

HSCT-derived epithelia showed significantly decreased TEER and MCC, indicating reduced epithelial barrier integrity and mucociliary function. For some HSCT-derived HAE as well as in the corresponding patients’ biopsies, markedly decreased differentiation was observed with loss of ciliated and mucus cells. These functional defects observed occurred in the context of increased expression of the epithelial progenitor cell markers, including KRT5, TP63 and SCGB1A1 and MUC5AC, a marker of goblet cells compared to LT-derived HAE, while an opposite trend for MKI67, a proliferation marker. This pattern may reflect activation of epithelial progenitor cells together with expansion of the secretory populations in response to HSCT-related injury. Increased KRT5 expression may indicate impaired progression toward terminal differentiation, as described in chronic airway injury^22^. In contrast, elevated MUC5AC and SCGB1A1 expression may indicate remodelling of secretory cell populations. Increased MUC5AC has been associated with mucus metaplasia rather than an increase in fully differentiated goblet cells^23^, while SCGB1A1 upregulation may indicate alterations in the club cell population involved in epithelial repair and regeneration^24,25^.

In contrast, markers associated with proliferation and multiciliate differentiation (MKi67, FOXJ1 and TFF3) did not show significant differences across groups, although directional trends were observed for some markers.

Reduced ciliation, impaired MCC and decreased TEER in some HSCT-derived epithelia are consistent with previous sinonasal epithelial studies reporting loss of ciliated and goblet cells, along with increased apoptosis in transplant recipients^26,27^. Alteration in cilia-related markers may also reflect impaired ciliogenesis or ciliary function as previously described^28,29^.

Although biopsies did not strictly sample the small airways, current evidence suggests that OB reflects a pan-bronchial or even a pan-respiratory process rather than a focal lesion^1^.

Within the HSCT cohort, some HAE showed near normal features, whereas others displayed more pronounced alteration. For example, HSCT3 exhibited marked changes across histological molecular and functional readouts and later developed BOS, whereas HSCT2 showed a similar histological alteration and comparable, albeit less pronounced, functional impairment, with subsequent recovery of lung function.

These observations illustrate the intra-cohort variability and limit direct extrapolation between epithelial phenotypes and clinical outcome.

#### LT epithelia

In contrast, LT-derived HAE displayed a more preserved phenotype and functional profile closer to NT controls, as reflected by TEER values and, to a lesser extent MCC measurements. A trend toward reduced MCC persisted and may be related to the decreased expression of DNALI1, a component of the axonemal dynein machinery required for ciliary motility, consistent with *in vivo* reports of early impaired mucociliary clearance^30,31^. Although LT epithelium has been exposed *in vivo* to chronic inflammatory, immune and mechanical constraints, these influences are no longer present after *ex vivo* reconstitution. The model therefore allows assessment of intrinsic and persistent epithelial features, partly independent of continuous external pressures. In this context, the relatively preserved functional readouts observed in LT-derived HAE may reflect stable post-transplant remodelling at a late stage in our cohort (mean 425 days post-transplantation), rather than complete epithelial normalization^32,33^. This is consistent with airway epithelial regeneration after lung transplantation, characterized by an early phase of injury and rapid repair, followed by a long-term remodelling phase associated with persistent alterations in some patients^34^. Mucin secretion was elevated compared with other groups. Although direct evidence for mucin hypersecretion after LT remains limited, this finding is consistent with reports of impaired MCC and mucus plugs in transplant recipients, suggesting persistent alterations in airway mucus homeostasis.

The different functional perturbations observed between LT and HSCT derived HAE suggest different patterns of airway epithelial remodelling after transplantation. HSCT- derived HAE were characterized by reduced TEER and MCC together with increased KRT5 expression, consistent with less differentiated phenotype. In contrast, LT-derived HAE displayed relatively preserved epithelial function but altered mucus homeostasis, with increased mucin secretion despite reduced MUC5AC expression. This discrepancy may reflect post-transcriptional regulation of mucin production or altered mucus release rather than increased goblet cell abundance. Both groups exhibited decreased DNALI1 expression compared to NT, suggesting that impaired terminal differentiation of the ciliated lineage may represent a common feature of post-transplant airway epithelial remodelling.

#### The ex vivo model

To our knowledge, this is the first study with fully ex vivo reconstituted HAE from bronchial epithelium *collected* from both LT and HSCT recipients using air-liquid interface culture, enabling detailed phenotypic characterization in a relevant system^35,36^. These models reproduce epithelial features observed *in vivo* and allow integrated functional and molecular analysis^37,38^. Comparison with FFPE biopsies reveals concordant epithelial phenotypes, supporting the relevance of this approach as a complementary model to investigate epithelial heterogeneity after transplantation.

Compared with short-term culture or *in vivo* tissue analysis, this system provides a standardized platform to assess epithelial properties independently of immune and endothelial influences. It also complements existing models and partially overcomes limitations of murine models, which lack distal bronchioles^35^.

Although small airways were not directly sampled, the concept of “United Airways” suggests a continuum between upper and lower airway epithelia, indicating that proximal airway alterations may reflect broader epithelial dysfunction^39^.

#### Epithelial chimerism

Beyond these functional and phenotypic differences, suggesting persistent transplantation-associated changes in epithelial biolofy, epithelial cellular origin may represent an additional layer contributing to the heterogeneity across transplant settings.

Epithelial chimerism has been previously reported^40^, but this is the first demonstration that chimeric epithelial cells can be maintained in *ex vivo* cultures, providing a valuable model for further characterization and investigation of its functional significance. In HSCT recipients, identification of epithelial cells of donor origin demonstrates that transplanted hematopoietic stem cells can differentiate into respiratory epithelial cells, which may have therapeutic implications. However, the low proportion of chimeric cells, even long after transplantation, raises questions about their actual contribution to repair. The timing of sampling suggests that donor-derived progenitors contribute to ongoing epithelial turnover rather than reflecting passive cell transfer^41^. In the most severely impaired sample (HSCT3), minimal chimerism was observed suggesting that donor-derived cells may not be actively recruited during extensive epithelial injury. Understanding the crosstalk between donor- and recipient-derived cells remain an important open question. Sequential HAE cultures could provide a unique platform to investigate these dynamics over time.

<u>Several limitations</u> should be considered. First, the sample size was modest. Nevertheless, the most severely impaired HSCT HAE corresponded to the patient who subsequently developed BOS within three months, providing preliminary support for the clinical relevance of the observed epithelial alterations. Second, LT and HSCT recipients were enrolled at different clinical stages, potentially introducing selection bias. HSCT recipients were included following a ≥10% FEV1 decline, whereas LT samples were obtained during routine follow up without FEV1-based criteria. Whether FEV1 changes in HSCT reflected pathology or measurement viability is uncertain, as respiratory function returns to pretransplant levels in all but one patient. Third, mucus characterization remains incomplete and warrants further investigations. Quantitative and qualitative mucus alterations could impair MCC, promote microbial persistence and contribute to airway obstruction, potentially influencing OB development. Fourth, epithelial chimerism was assessed using RNA-FISH on *ex vivo* HAE from sex mismatched LT and HSCT recipients, and SNP-based analysis on all HAE from all HSCT. SNP-based approaches could not be applied to FFPE biopsy, due to potential false positive from donor blood cells which are excluded during *ex vivo* HAE reconstitution. This analysis cwas not feasible in LT recipients because donor genomic sequences were unavailable. Although, the correlation observed in the HSCT sample points a potential relationship, its broader applicability remains to be confirmed.

Finally, the model excludes immune and endothelial components, omitting the study of epithelial and immune cells. Nevertheless, it provides a controlled system for epithelial specific analysis and functional readouts.

## Conclusion

We report the first fully differentiated HAE model derived from HSCT and LT recipients, allowing direct comparison with NT controls. HSCT epithelia showed heterogeneous and occasionally impaired differentiation and function, whereas LT epithelia largely resembled NT epithelia, highlighting transplant procedure-specific epithelial remodelling. The model also revealed epithelial chimerism, reflecting the contribution of progenitor cells derived from the donor in HSCT recipients and from the recipient in LT recipients. Despite modest sample size and the absence of immune blood cells and endothelial components, this *ex vivo* platform provides a physiologically relevant tool to explore airway epithelial dynamics, remodelling and mechanisms underlying post-transplant airway disease.

## Acknowledgement

The patient cohort used in this study overlaps with that used in a separate manuscript from the same authors; the present study addresses distinct research questions and reports different findings^42^.

The authors thank the Department of Genetic Medicine, Laboratory and Pathology, University Hospitals of Geneva (HUG) and the Cell Therapy and Transplantation Laboratories team at the University Hospital of Geneva for their technical assistance. The authors also thank Doctor Delphine Gras and Professor Pascal Chanez for sharing their expertise.

## Disclosures

The authors declare that they have no relevant conflicts of interest.

This work was supported by the Fondation Privée des Hôpitaux Universitaires de Genève (HUG), the Fondation La Laurène, Benoît Ficheur and OrganoVIR Labs. The authors declare that AI or AI-assisted technologies were not used to generate the scientific content of the manuscript.

## Authorship Contributions

The study was conceived and designed by AB, CT and LB. AB, CT and LB wrote the manuscript. All authors (JS, SL, CDV, CG, YC, FG, GB, RM, JLG, JV, AB, LB and CT) critically revised the manuscript for important intellectual content and approved the final version. All authors agree to be accountable for all aspects of the work, ensuring that questions related to accuracy or integrity are appropriately investigated and resolved.

## Ethics approval

The project received the approval of the Swiss Ethic Committee. Project number 2023-00140 approved on March 7^th^, 2023.

## Notes

### Competing Interest Statement

The authors have declared no competing interest.

