## supplementary materials for "Ex vivo human airway epithelia modelling reveals specific alterations in lung and hematopoietic stem cell transplant recipients"

** Co last authors*

**Methods**

<u>Presence of epithelial chimerism in reconstituted human airway epithelia</u>

*Fluorescence in situ hybridization (FISH):*

Chimerism in airway epithelial cells from sex-mismatched transplant recipients was assessed by FISH using X and Y chromosome-specific probes. During the study. Over the course of the study, the procedure evolved from a manual workflow to a semi-automated approach. Both approaches relied on the same reagents, hybridization conditions and analysis criteria.

Formalin-fixed paraffin-embedded (FFPE) or cytospin preparation of airway epithelial cells from *ex vivo* reconstituted HAE were processed for FISH. Cells were hybridized with commercially available centromeric X (orange spectrum) and Y (green spectrum) probes (Abbot, 30-171050) or ZytoLight CEN X/Y Dual Color Probe, ZYtoVision (Z-2120-200), according to the manufacturer's instructions.

For the semi-automated workflow, staining was performed on the Discovery Ultra platform (Ventana/Roche). After rehydratation, samples were treated with ISH protease 3 (Ventana, 780-4149) for 12 minutes at 37°C, followed by fixation in 4& formaldehyde for 4 minutes. Slides were then dehydrated, airdried and 10 µl of XY probes were applied before cover slipping and sealing. Denaturation was carried out at 79°C for 8 minutes, followed by hybridization at 37°C for 20h. Post-hybridization washes were performed at 80°C in 2x SSC buffer. Slides were subsequently dehydrated and mounted with a DAPI-containing medium (DAPI Counterstain II, Abbott, 30-804930).

For the manual workflow, hybridization and post-hybridization steps were performed according to the same conditions above.

Fluorescent signals were visualized using a Zeiss Axio Imager microscope equipped with appropriate filter sets.

Image acquisition was performed using the PhenoImager HT system (Akoya Biosciences) with a 40× objective, corresponding to a pixel resolution of 0.25 µm.

*Next-generation sequencing (NGS) based assay:*

Chimerism was quantified using the AlloSeq HCT platform (CareDx, Brisbane, CA, USA), a targeted next-generation sequencing-based assay enabling sensitive discrimination of the DNA from the donor and the recipient. Genomic DNA was extracted from epithelial cell fractions and analysed using a multiplex panel of 202 highly polymorphic single-nucleotide polymorphism (SNP) markers distributed across multiple autosomal chromosomes. These SNPs were selected for high minor allele frequency and limited genetic linkage, increasing the likelihood of informative differences between donor and recipient genotypes. Target regions were amplified by multiplex PCR, followed by indexed library preparation and sequencing on an Illumina platform according to the manufacturer's protocol. Informative loci were defined using pre-transplant donor and recipient genotypes, and allele frequencies were quantified with AlloSeq HSCT analysis software (version 2-2-0) to estimate the proportion of donor-derived DNA in epithelial samples. The approach allows quantitative chimerism measurement with a sensitivity of approximately 0.1-0.3%, depending on the number of informative markers available.

**Table S1 – Clinical characteristics of non-transplant control patients**

| | NT1 | NT2 | NT3 | NT4 | NT5 | NT6 |
|---|---|---|---|---|---|---|
| **Age (years), sex** | 67, F | 50, M | 67, M | 74, M | 62, F | 60, M |
| **Smoking history (pack-years)** | 40 | 20 | 20 | No | No | 40 |
| **Indication for bronchoscopy** | Mediastinal lymph node | COPD nodule | Nodule | Nodule with adenopathy | Mediastinal lymph node | Nodule |
| **FEV1 (ml, % predicted)** | 1610 (71) | 1610 (47) | 2330 (89) | 2100 (82) | 2510 (83) | 1560 (48) |
| **FVC (ml, % predicted)** | 2460 (85) | 2200 (51) | 3460 (102) | 2620 (82) | 3170 (86) | 2130 (51) |
| **FEV1/FVC ratio** | 0.65 | 0.73 | 0.67 | 0.80 | 0.79 | 0.73 |

*NT: non-transplant; pack-year (number of cigarette packs per day x the number of years of smoking) FEV1: forced expiratory volume in 1 second, FVC: forced vital capacity.*

**Table S2 – Clinical characteristics of LT recipients**

| | LT1 | LT2 | LT3 | LT4 | LT5 | LT6 |
|---|---|---|---|---|---|---|
| **Sex mismatch** | No | Yes | NA | Yes | No | No |
| **Recipient smoking history (pack-years)** | 30 | 0 | 40 | 25 | 10 | 10 |
| **Donor age (years)** | 22 | 71 | NA | 36 | 46 | 52 |
| **Donor smoking history (pack-years)** | 4 | 10 | NA | 0 | 0 | 0 |
| **Indication for transplantation** | COPD | Emphysema | IPF | IPF /PAH | IPF | NSIP |
| **Type of transplantation** | Bilateral | Bilateral | Bilateral | Bilateral | Bilateral | Bilateral |
| **Serology (Recipient/Donor)** | CMV +/- EBV +/+ | CMV +/+ EBV +/+ | CMV -/- EBV+/+ | CMV +/- EBV+/- | CMV +/- EBV +/+ | CMV +/+ EBV +/- |
| **Blood Group (Recipient/Donor)** | A +/A + | A+/A+ | B+/NA | A+/A+ | O+/O+ | O+/O- |
| **Primary graft dysfunction** | No | No | No | Yes (grade 3) | Yes (grade 3) | Yes (grade 3) |

| | | | | | | |
|---|---|---|---|---|---|---|
| **Days post-transplant to biopsy** | 725 | 574 | 545 | 276 | 96 | 334 |
| **Ischemic time (h:min)** | 5:37 | 4:58 | NA | 10:00 | 4:53 | 9:45 |
| **Gastroesophageal reflux** | Yes | No | Yes | No | No | Yes |
| **Respiratory viral infection post-transplant (Type, days)** | RHV (477) | SARS-CoV-2 (67) | SARS-CoV-2 (511) | None | None | RHV (-44); RSV (-107) |
| **Respiratory bacterial infection post-transplant (Type, days)** | None | *Stenotrophomonas spp.* (477) | *Pseudomonas aeruginosa* (311) | None | None | *Corynebacterium spp.* (483) |
| **Respiratory fungal infection post-transplant (Type, days)** | *Aspergillus terreus* (182) | None | *Aspergillus versicolor* (360) | *Candida albicans* (64) | *Aspergillus niger* (25) | None |
| **Immunosuppressive therapy at biopsy** | Tacrolimus 3.75mg BID; MMF 1g BID; Prednisone 5mg | Tacrolimus 3mg BID; MMF 180mg BID; Prednisone 20mg | Tacrolimus 4.5mg BID; MMF 360mg BID; Prednisone 7.5mg | Tacrolimus 2.75mg BID; MMF 500mg BID; Prednisone 5mg | Tacrolimus 2.5mg BID; MMF 1g BID; Prednisone 22.5mg | Tacrolimus 2.5mg BID; MMF 500mg BID; Prednisone 12.5mg |

| | | | | | | |
|---|---|---|---|---|---|---|
| **FEV1 (ml, % predicted)** | 2560 (110) | 2170 (76) | 3540 (103) | 2760 (103) | 2780 (87) | 1690 (63) |
| **FVC (ml, % predicted)** | 2670 (92) | 2170 (60) | 4160 (90) | 3690 (110) | 3800 (92) | 2850 (85) |
| **FEV1/FVC ratio** | 0.96 | 1 | 0.85 | 0.75 | 0.73 | 0.77 |

*LT : lung transplantation; pack-year (number of cigarette packs per day x the number of years of smoking); IPF: idiopathic pulmonary fibrosis; COPD: chronic obstructive disease; PAH: pulmonary hypertension; NSIP: non-specific interstitial pneumonia; CMV: cytomegalovirus; EBV: Epstein Barr virus; Toxo: toxoplasmosis; HPIV3: human parainfluenza virus type 3; RHV: rhinovirus; RSV: respiratory syncytial virus; MMF: mycophenolate mofetil; BID: bis in die (i.e. twice daily); TID : ter in die (i.e. three times daily); FEV1: forced expiratory volume in 1 second, FVC: forced vital capacity.*

**Table S3 – Clinical characteristics of HSCT recipients**

| Parameter | HSCT1 | HSCT2 | HSCT3 | HSCT4 | HSCT5 | HSCT6 |
|---|---|---|---|---|---|---|
| Age (years), sex | 61, M | 28, F | 59, F | 47, M | 30, M | 55, F |
| Donor age (years), sex | 52, F | 57, F | 32, F | 23, F | 32, M | 65, F |
| Smoking history (pack-years) | 20 | 0 | 26 | 20 | 0 | 0 |
| Indication for HSCT | CLL | AML | NHL T | ALL | Sickle cell Disease | ALL |
| Graft type | Haploidentical, PBSC, T-cell replete | Matched unrelated donor (10/10), PBSC, T-cell replete | Haploidentical, PBSC, T-cell replete | Haploidentical, PBSC, T-cell replete | Haploidentical, bone marrow, T-cell replete | Matched unrelated donor (10/10), PBSC, T-cell replete |
| GVHD prophylaxis | Cyclosporin A 3mg/kg/d MTX 15mg/m2 | Tacrolimus 0.03mg/kg/d MTX 15mg/m2 | Cyclophosphamide 50mg/kg/d Tacrolimus | Cyclophosphamide 50mg/kg/d Tacrolimus | Cyclophosphamide 50mg/kg/d Tacrolimus | Tacrolimus 0.03mg/kg/d MTX 15mg/m2 |

| | | | | | | |
|---|---|---|---|---|---|---|
| | | | 0.03mg/kg/d MMF 15mg/kg TID | 0.03mg/kg/d MMF 15mg/kg TID | 0.03mg/kg/d MMF 15mg/kg TID | |
| **Days post-transplant to biopsy** | 892 | 210 | 343 | 1008 | 120 | 359 |
| **Serology (Recipient/Donor)** | CMV -/- Toxo -/- EBV -/+ | CMV +/+ Toxo +/- EBV +/+ | CMV -/- Toxo +/+ EBV+/+ | CMV +/+ Toxo +/- EBV+/- | CMV -/- Toxo -/- EBV +/+ | CMV +/+ Toxo +/+ EBV -/+ |
| **Donor blood chimerism at biopsy (%)** | 100% | 93% | 100% | 100% | 99% | 99% |
| **Conditioning regimen (total dose)** | MAC ATG (1.5g) Cyclophosphamide (5.6g) TBI (12Gy) | MAC ATG (2400mg) Fludarabine (275mg) Treosulfan (77g) | RIC Clofarabine (270mg) Cyclophosphamide (2.3g) Fludarabine (270mg) | MAC ATG (700mg) Etoposide (4.2g) TBI (10Gy) | RIC ATG (300mg) Thiotepa (651mg) Cyclophosphamide (1822mg) Fludarabine (257mg) TBI (2Gy) | RIC ATG (500mg) Fludarabine (275mg) TBI (8Gy) |

| | | | | | | |
|---|---|---|---|---|---|---|
| | | | Melphalan (198mg) | | | |
| **History of GVHD** | Grade 3 acute (skin) | Mild chronic (skin) | Grade 2 acute (gastrointestinal) | Mild chronic (skin) | Grade 2 acute (gastrointestinal) | Grade 3 acute (gastrointestinal) |
| **GVHD at the time of biopsy** | Mild chronic (skin) | None | Severe chronic (skin) | None | Mild chronic (skin) | None |
| **Respiratory viral infection post-transplant (Type, days)** | None | MPV; SARS-CoV-2; RHV (441) | RHV (151, 263); SARS-CoV-2 (753) | HPIV3; RHV (83) | RHV (-134); HPIV3 (156, 230); RHV; Cor-OC43 (267); RSV (355) | None |
| **Respiratory bacterial infection post-transplant (Type, days)** | None | None | None | None | None | None |

| | | | | | | |
|---|---|---|---|---|---|---|
| **Respiratory fungal infection post-transplant (Type, days)** | None | None | None | None | *Aspergillus niger* (60) | None |
| **Immunosuppressive therapy at biopsy** | MMF 750mg BID Prednisone 20mg Ruxolitinib 5mg BID | Azacitidine | Prednisone 10mg Tacrolimus 2.5mg BID | None | Prednisone 5mg Tacrolimus 2.5mg | Ponatinib 15mg/j |
| **FEV1 (ml, % predicted)** | 2750 (77) | 1710 (77) | 2540 (78) | 3760 (97) | 1900 (63) | 1760 (73) |
| **FVC (ml, % predicted)** | 3280 (80) | 2110 (76) | 2910 (76) | 4360 (90) | 2050 (58) | 2480 (81) |
| **FEV1/FVC ratio** | 0.84 | 0.81 | 0.87 | 0.86 | 0.93 | 0.71 |

*HSCT: hematopoietic stem cell transplantation; pack-year (number of cigarette packs per day x the number of years of smoking); PBSC: peripheral blood stem cell; HLA: human leukocyte antigen; AML : acute myeloid leukaemia; ALL: acute lymphoid leukaemia; MTX: methotrexate; MMF: mycophenolate mofetil; CMV: cytomegalovirus; EBV: Epstein Barr virus; Toxo: toxoplasmosis; MAC: myeloablative conditioning; RIC:*

*reduced-intensity conditioning; ATG: anti thymoglobulin, TBI: total body irradiation; GVHD: graft versus host disease; HPIV3: human parainfluenza virus type 3; RHV: rhinovirus; RSV: respiratory syncytial virus; MPV: metapneumovirus; CoroOC43: coronavirus OC43; BID: bis in die (i.e. twice daily); FEV1: forced expiratory volume in 1 second, FVC: forced vital capacity.*

Table S4 – Individual functional parameters from NT-derived HAE

| Parameter | NT1 | NT2 | NT3 | NT4 | NT5 | NT6 |
|---|---|---|---|---|---|---|
| **TEER ($\Omega.cm^2$)** | 894 | 754 | 824 | 736 | 815 | 796 |
| **MCC (**µm/s) | 64.6456 | 46.80356 | 38.12233 | 17.80279 | 46.43061 | 44.69308 |
| **Mucin (OD)** | 0.28183824 | 0.29109087 | 0.45834139 | 0.4015 | 0.38785 | 0.34475 |

*TEER: transepithelial electrical resistance assessing barrier integrity; MCC: Mucociliary clearance, quantified as microbeads velocity across HAE apical surface; Mucin quantified by ELLA assay*

Table S5 – Individual functional parameters from HSCT-derived HAE

| Parameter | HSCT1 | HSCT2 | HSCT3 | HSCT4 | HSCT5 | HSCT6 |
|---|---|---|---|---|---|---|
| **TEER ($\Omega.cm^2$)** | 483 | 597 | 252 | 535 | 513 | 521 |

| | | | | | | |
|---|---|---|---|---|---|---|
| **MCC** (µm/s) | 27.59229 | 13.05379 | 5.791107 | 41.7163 | 13.65023 | 39.92064 |
| **Mucin (OD)** | 0.41625 | 0.38480639 | 0.27652012 | 0.55525 | 0.45834139 | 0.58305 |

*TEER: transepithelial electrical resistance assessing barrier integrity; MCC: Mucociliary clearance, quantified as microbeads velocity across HAE apical surface; Mucin quantified by ELLA assay*

Table S6 – Individual functional parameters from LT-derived HAE

| **Parameter** | **LT1** | **LT2** | **LT3** | **LT4** | **LT5** | **LT6** |
|---|---|---|---|---|---|---|
| **TEER (Ω.cm²)** | 874 | 755 | 807 | 723 | 750 | 812 |
| **MCC** (µm/s) | 31.95782 | 41.09624 | 31.92778 | 28.32153 | 33.45851 | 39.02331 |
| **Mucin (OD)** | 0.5974 | 0.42564341 | 0.53845 | 0.63915 | 0.61837903 | 0.56571431 |

*TEER: transepithelial electrical resistance assessing barrier integrity; MCC: Mucociliary clearance, quantified as microbeads velocity across HAE apical surface; Mucin quantified by ELLA assay*

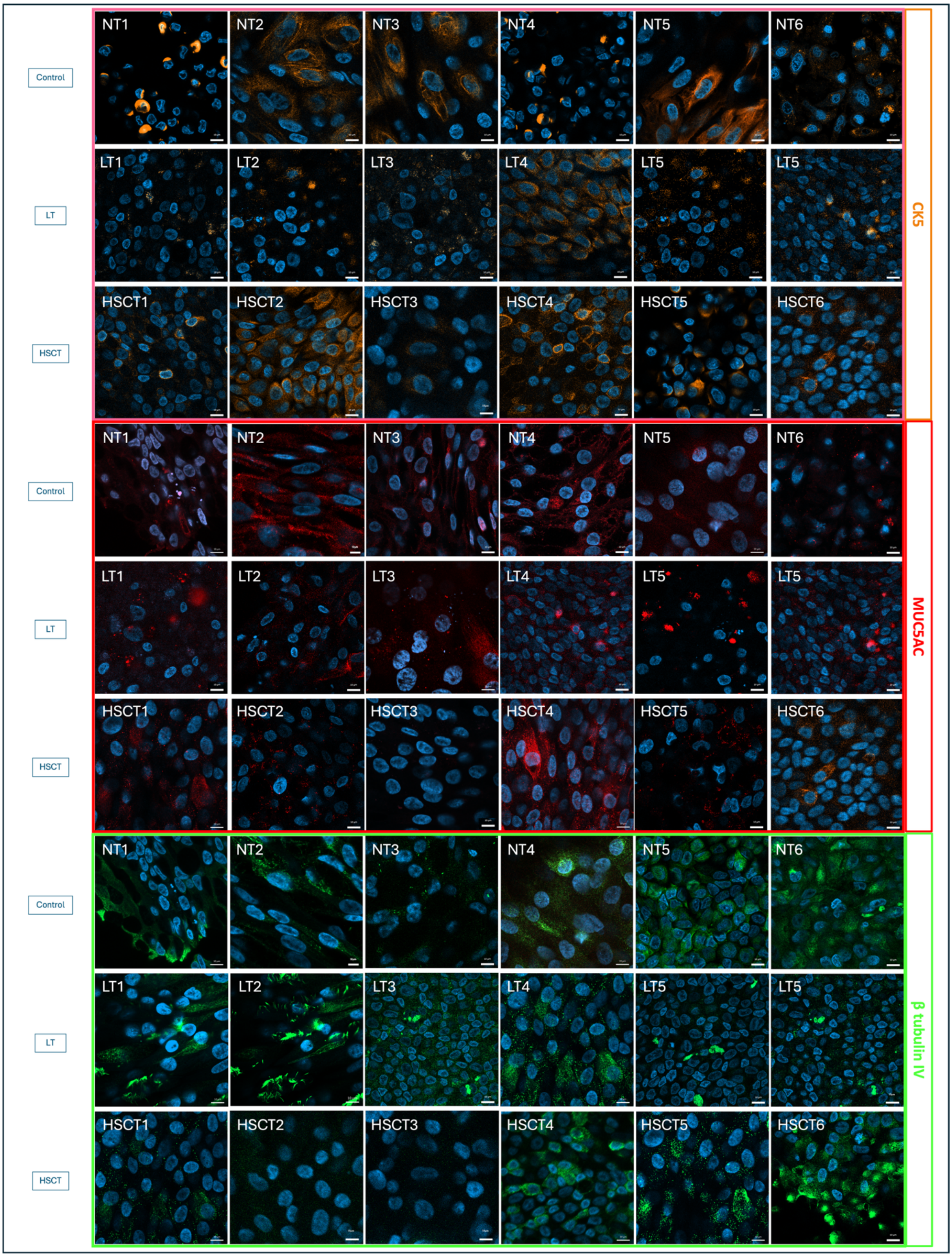


**Figure S1- Immunofluorescence of human airway epithelium from all donors**

*Immunofluorescence of fully differentiation HAE at day 30 of culture in transwell. The figure is composed of three panels (A-C), each arrange in three rows representing the three donor groups: NT, LT and HSCT; and six columns corresponding to six individual donors per group. (A) Two-dimensional view of HAE showing basal cells stained for cytokeratin 5 (CK5, orange); (B) Goblet cells stained for MUC5AC in red; (C) Ciliated cells stained for β-tubulin IV in green. Nuclei were counterstained with DAPI (4'-6-Diamidino-2-phenylindole dihydrochloride). Images were acquired using confocal microscopy (Axio Imager, LSM800) with a 63x objective, scale bars: 10μm*

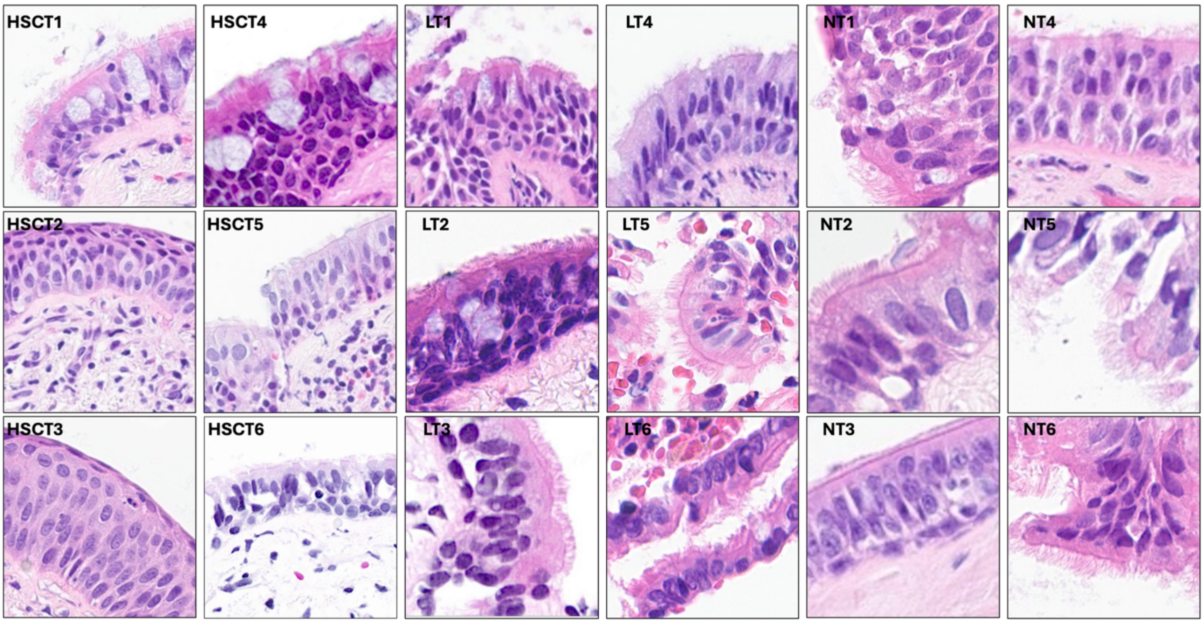


**Figure S2 -Paraffin-embedded sections prepared from bronchial biopsies**

*Biopsies obtained from 6 HSCT recipients; 6 LT recipients and 6 controls were stained with Haematoxylin and eosin stain (H&E stain). The bronchial epithelium presented an aspect of metaplasia with fewer ciliated and mucous cells in biopsies from HSCT recipients 2 and 3 and few mucous cells in HSCT5. Bronchial epithelium of HSCT1, 4, and 6 showed pseudo-stratified aspect made of ciliated, mucous, and basal cells similarly to that of LT recipients and controls.*

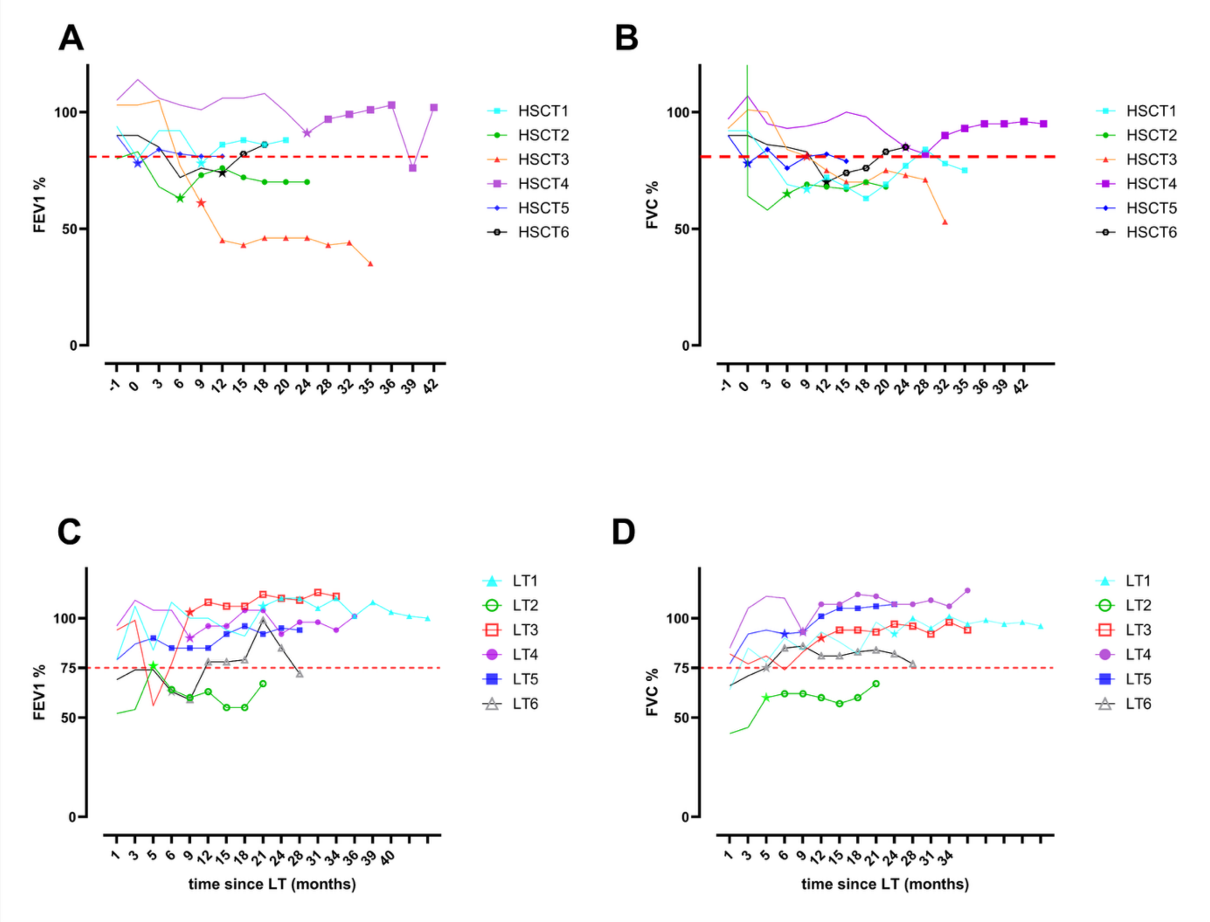


**Figure S3- Lung function trajectory before and after biopsy of transplant recipients**

*(A) FEV1 trajectory of HSCT recipients, showing a significant and persistent FEV1 decline in HSCT3. (B) FVC trajectory of HSCT recipients. (C) FEV1 trajectory of lung transplant recipients (D)FVC trajectory of lung transplant recipients. Trajectories are represented before biopsy (continuous line), at the time of biopsy (star) and after biopsy (dots). FEV1: forced expiratory volume in 1 second, FVC: forced vital capacity.*
